# Targeted analysis of dyslexia-associated regions on chromosomes 6, 12 and 15 in large multigenerational cohorts

**DOI:** 10.1101/2023.08.01.551585

**Authors:** Nicola H. Chapman, Patrick Navas, Michael O. Dorschner, Michele Mehaffey, Karen G. Wigg, Kaitlyn M. Price, Oxana Y. Naumova, Elizabeth N. Kerr, Sharon L. Guger, Maureen W. Lovett, Elena L. Grigorenko, Virginia Berninger, Cathy L. Barr, Ellen M. Wijsman, Wendy H. Raskind

## Abstract

Dyslexia is a common specific learning disability with a strong genetic basis that affects word reading and spelling. An increasing list of loci and genes have been implicated, but analyses to-date investigated only limited genomic variation within each locus with no confirmed pathogenic variants. In a collection of >2000 participants in families enrolled at three independent sites, we performed targeted capture and comprehensive sequencing of all exons and some regulatory elements of five candidate dyslexia risk genes (*DNAAF4*, *CYP19A1*, *DCDC2*, *KIAA0319* and *GRIN2B*) for which prior evidence of association exists from more than one sample. For each of six dyslexia-related phenotypes we used both individual-single nucleotide polymorphism (SNP) and aggregate testing of multiple SNPs to evaluate evidence for association. We detected no promoter alterations and few potentially deleterious variants in the coding exons, none of which showed evidence of association with any phenotype. All genes except *DNAAF4* provided evidence of association, corrected for the number of genes, for multiple non-coding variants with one or more phenotypes. Results for a variant in the downstream region of *CYP19A1* and a haplotype in *DCDC2* yielded particularly strong statistical significance for association. This haplotype and another in *DCDC2* affected performance of real word reading in opposite directions. In *KIAA0319*, two missense variants annotated as tolerated/benign associated with poor performance on spelling. Ten non-coding SNPs likely affect transcription factor binding. Findings were similar regardless of whether phenotypes were adjusted for verbal IQ. Our findings from this large-scale sequencing study complement those from genome-wide association studies (GWAS), argue strongly against the causative involvement of large-effect coding variants in these five candidate genes, support an oligogenic etiology, and suggest a role of transcriptional regulation.

**Author Summary:** Family studies show that genes play a role in dyslexia and a small number of genomic regions have been implicated to date. However, it has proven difficult to identify the specific genetic variants in those regions that affect reading ability by using indirect measures of association with evenly spaced polymorphisms chosen without regard to likely function. Here, we use recent advances in DNA sequencing to examine more comprehensively the role of genetic variants in five previously nominated candidate dyslexia risk genes on several dyslexia-related traits. Our analysis of more than 2000 participants in families with dyslexia provides strong evidence for a contribution to dyslexia risk for the non-protein coding genetic variant rs9930506 in the *CYP19A1* gene on chromosome 15 and excludes the *DNAAF4* gene on the same chromosome. We identified other putative causal variants in genes *DCDC2* and *KIAA0319* on chromosome 6 and *GRIN2B* on chromosome 12. Further studies of these DNA variants, all of which were non-coding, may point to new biological pathways that affect susceptibility to dyslexia. These findings are important because they implicate regulatory variation in this complex trait that affects ability of individuals to effectively participate in our increasingly informatic world.

## INTRODUCTION

Dyslexia is a complex disorder of neurobiological origin that can be defined as unexpectedly low accuracy and/or rate of oral reading of single words or pronounceable pseudowords, or low accuracy of spelling [1]. It manifests as difficulty in learning to read and spell despite adequate instruction and is not attributable to general cognitive impairment, primary sensory or motor impairment, psychiatric or other neurologic disorder or delays in aural or oral language. The stated prevalence of dyslexia varies depending on ascertainment schemes, exclusion criteria, tests included in diagnostic assessment, and thresholds used for a categorical diagnosis. In school-aged children, most estimates of dyslexia fall between 5-12% [2–5] but have been as low as 3.5% [6] and as high as 20% [7]. In almost all past studies, including our own, males are at greater risk than females for both presence of dyslexia and its severity [2, 5, 8–10]. Even with educational intervention, many aspects of dyslexia can persist into adulthood, including slow reading speed and poor spelling-related writing abilities [11–13] leaving lasting impacts on self-esteem, educational opportunities, and occupational choices [14–16].

Multiple lines of evidence, including twin [17], familial aggregation [8, 18], adoption [19], and linkage and/or association studies [20], have led to the consensus that there is a substantial genetic contribution to dyslexia and its component phenotypes. Heritability estimates are as high as 50-70% [21, 22]. Although rare families have been described in which dyslexia appears to be transmitted as a single gene disorder [23–28] studies in the general population show that, like most complex traits, dyslexia and its correlated underlying processes are genetically heterogeneous and likely involve the influence of variation in multiple genes [29]. Such heterogeneity complicates identification of underlying genes, regardless of the study design, but multiple candidate susceptibility genes have been nominated from genomic regions of interest (ROIs) identified by linkage analyses [30, 31], genome wide association studies, GWAS, [32–34], copy number scan [35], structural chromosome rearrangements [36, 37], or whole genome sequencing [27]

Of the multiple reported ROIs for dyslexia risk, a small number have the strongest support. DYX1 on chromosome 15q [38–44] and DYX2 on chromosome 6p [39, 45–48] are the loci with the greatest replication across samples. The University of Washington (UW) and Hospital for Sick Children (SickKids, SK) samples have implicated several additional ROIs, with one of the most convincing on chromosome 12q [49]. Candidate genes that have been most studied in the ROIs on chromosomes 15q and 6p include Dynein Axonemal Assembly Factor 4 (*DNAAF4*, previously named *DYX1C1*/*EKN1*) on chromosome 15q [50–52], and Double Cortin Domain Containing 2 (*DCDC2*) and *KIAA0319* on chromosome 6p [53–61]. Support for involvement of these and other candidate risk genes has been reported from a variety of association and linkage studies in humans, and functional studies of brain development in rodents, but evidence is inconsistent [62–68]. There have been failures to detect linkage [69–71] or association [72–78], reports of association to opposite alleles [51, 52, 72], and demonstration of functional competence of the putative risk allele [79]. Meta-analyses have not resolved these conflicts [76, 80–82]. Nor have GWAS, which have provided at most weak support for the loci [34, 83–88].

This includes a recent large GWAS that did not detect significant evidence of association with any reported candidate dyslexia risk gene [32]. Variability in conclusions across the different study designs is not surprising: genome-wide linkage analyses and GWAS both allow location scans, but linkage analyses are most sensitive to relatively rare variation of large effect and GWAS are most sensitive to common, smaller-effect variation. Neither approach queries all the DNA variation, which requires more-expensive DNA sequencing of (at least) the regions of interest.

The putative effect of candidate genes on neuronal migration has been used to bolster their credibility [89, 90], given early reports of cortical brain abnormalities in people who were thought to have had dyslexia [91, 92]. However, although cortical abnormalities have been observed with knockdown of the rat orthologues of *Dnaaf4* [79], *Kiaa0319* [62], or *Dcdc2* [93], this is not observed in knockout mice [94–96], and the cortical migration hypothesis remains unproven [65]. Observations that dyslexia candidate genes seem to have a role in ciliogenesis [97, 98], synaptic transmission [99], or axonal growth [100], have led to alternative hypotheses of pathogenesis.

Although such issues are to be expected in a complex disorder, to date, no causative pathogenic variants have been confirmed. Some possible explanations for this failure include: (1) genetic and/or phenotypic heterogeneity that masks detection in samples ascertained and phenotyped with different criteria; (2) the risk element(s) may alter expression of the protein but not its amino acid sequence; (3) the risk element may escape recognition, but affect splicing; and (4) the number of samples sequenced comprehensively has been too small to have the power to detect variants of modest effect size.

Recent advancements in DNA sequencing methods now enable the larger scale sequencing efforts that are necessary to evaluate genetic variation in ROIs more comprehensively than was possible earlier. This technology allows us, in this current multi-site study, to investigate the potential role of variants of smaller effect size, non-coding variants, and sample heterogeneity as possible explanations for previous variable results in ROIs implicated in dyslexia. To search for variants that show evidence of association with dyslexia phenotypes, we report the results of genomic sequencing and association analyses in a collection of >2000 participants in families with dyslexia and shared phenotypic measures enrolled at three institutions. We carried out a comprehensive analysis of the coding region and some regulatory element motifs of five putative dyslexia risk genes and found additional evidence for a role in dyslexia risk for *DCDC2*, *KIAA0319*, *GRIN2B* and *CYP19A1*, but not for *DNAAF4*.

## METHODS

### Participants and Phenotypes Across Sites

#### Sample and phenotype selection strategy

We selected participants enrolled in studies of dyslexia at three institutions that had overlapping phenotype batteries: University of Washington (UW), The Hospital for Sick Children (SickKids; SK), and the University of Houston (UH). Phenotypes were measured by standardized normed tests administered by more than one site. This strategy provided a large total number of participants screened with an essentially equivalent test-battery allowing for joint analysis across the three cohorts while minimizing introduction of excess phenotypic heterogeneity. The sample is largely separate from other cohorts analyzed for association of dyslexia with the ROIs investigated in the current study, thus providing an independent evaluation. Under the assumption that genetic heterogeneity in dyslexia may be reflected in phenotypic heterogeneity, the focus was on individual subtests or index scores based on multiple subtests in standardized, nationally normed tests that are predictive of reading or spelling outcomes or that assess processes related to reading and spelling achievement such as verbal reasoning or phonological memory. To maximize sample size, only test measures administered by more than one of the three cohort sites described below were evaluated. For all cohorts, individuals with evidence of intellectual disabilities, neurological or severe psychiatric disorder, or known genetic disorder associated with language impairment were excluded.

##### University of Washington (UW) Cohort

Recruitment and evaluation of probands and their multigenerational family members were done under a protocol approved by the Institutional Review Board of the UW and are described elsewhere in detail [8, 11] and briefly summarized in the Supporting Information document. The full data set consists of 2079 individuals in 284 families. Of the available subjects, 1347 individuals from 278 families provided quality DNA samples for sequencing. Phenotypic data were available for 96.8% of the 1333 samples that passed quality control (QC) testing. Family sizes ranged from 3 to 51 individuals, with a median family size of 12 in 2-4 generations. Self-reported ethnicities were as follows: non-Hispanic White (90%), Asian (2.1%), Native American (2.0%), African American (1.1%), Hispanic (0.8%), and Pacific Islander (0.2%).

##### The SickKids (SK) Cohort

Details of the ascertainment, assessment, and inclusion/exclusion criteria have been described previously [101, 102] and a brief description is provided in the Supportive Information document. Written informed consent and/or assent was obtained from all participants under protocols approved by the Hospital for Sick Children and University Health Network Research Ethics Boards. The SK cohort comprised 816 participants from 245 families (155 with the proband and one or both parents and 90 with two or more children and one or both parents). Phenotypes were available for 99% of children (349 of 351), and 85% of parents (394 of 465). The phenotype battery in parents was limited to TOWRE measures of single word reading efficiency and phonological decoding efficiency. Self-reported ancestry was available for 185 (76%) of the families. Of these, 180 (73.5%) reported European or European-Canadian ancestry. The remaining families reported small amounts of indigenous ancestry (3 families, 1.6%), African ancestry (1 individual) and Mexican ancestry (1 individual).

##### The University of Houston (UH) Cohort

The subset of families of probands with dyslexia used here is a component of the ongoing collection supported by the Florida Learning Disabilities Research Center (FLDRC). Written informed consent was obtained from all participants under protocols approved by the Research Ethics Boards of Yale University or the University of Houston. The probands were children aged 8–17 who had problems reading and were native English speakers. They were registered with FLDRC and recruited through their registration. The probands and their siblings in the same age range were administered a battery of tests including IQ and reading abilities. Designation of affected status required the Wechsler Abbreviated Scale of Intelligence, WASI [103], score ≥80 and a score at least 1.5 SD below the mean on 2 of 3 measures of single real- or non-word reading, or 1 SD below the mean on all 3 indicators. Self-reported ethnicities were as follows: African American (13.5%) and Hispanic (86.5%). This cohort consisted of 49 participants (37 children and 12 parents) from 15 nuclear families. Parents provided blood samples but were not phenotyped.

### Molecular Methods

#### Overview of approach

The targeted capture approach used here was designed to produce a comprehensive assessment of sequence variants relevant to the subset of dyslexia candidate genes evaluated. In the large, combined sample, this included all coding variants, as well as non-coding variants potentially involved in some aspects of gene regulation. For this investigation, the focus was on the potential impact on RNA processing and/or transcription-factor (TF) motifs in tissue-dependent open chromatin regions that control gene expression.

#### Genomic regions investigated

We targeted DNA sequence in three previously published regions of interest (ROIs) for comprehensive evaluation. Two of the ROIs have been widely investigated, with both positive linkage and association analyses reported by more than one group: the DYX1 locus on chromosome 15q and the DYX2 locus on chromosome 6p. Previous linkage analyses in the UW cohort provided support for DYX1 as a dyslexia candidate region [52, 70] and similar studies in subsets of the SK cohort supported both DYX1 and DYX2 [101, 104, 105]. From each of these ROIs, we selected for study two genes based on prior publications supporting a role in dyslexia: in DYX1, *DNAAF*4 [40, 44, 50–52] and C*YP19A1* [37, 106, 107]; and in DYX2, *KIAA0319* [54, 62, 77, 108, 109] and *DCDC2* [81, 110, 111]. Intron 2 of *DCDC2* [93] contains READ1 – a complex compound STR polymorphism that was previously proposed as the functional dyslexia-risk component in the DYX2 region [112–114] and was included as part of our capture-design. We also selected the glutamate ionotropic receptor NMDA type subunit 2B *(GRIN2B)*, in a third ROI, on chromosome 12p. This region was among those with the strongest evidence of linkage from the UW family studies, with support across several test-battery items [49, 115, 116]. *GRIN2B* has also been implicated as a dyslexia risk factor [117–119]. For analysis of these five genes, we developed a set of custom capture probes to enable comprehensive evaluation of potential regulatory and splice region sequence variants in addition to coding region variants, as described below.

#### Sample preparation

For most samples, genomic DNA was extracted from peripheral blood mononuclear cells or Epstein–Barr virus-transformed B-lymphoblastoid cell lines. When only saliva samples were available, DNA was extracted using a DNAGenotek OGR-500 kit (DNAGenotek Inc, Ontario, Canada) according to the manufacturer’s instructions.

#### Single molecule molecular inversion probe (smMIP) targeted capture and sequencing

The capture target consisted of all potentially functional sequences and variants in each ROI around and including each of the five selected genes. This included a total of 277 potential regulatory regions. We used smMIPs to capture targeted DNA with methods described elsewhere [120, 121], followed by multiplex sequencing. Additional details are provided in the Supporting Information document.

We used the UW pipeline [122] to design smMIPs to capture all exons and 10-20bp of flanking intron sequences. This approach ensured capture of the splice site branch A points (RefSeq, hg19/GrCH37 build [121]). All analysis was on this genome build. For the non-coding regulatory regions, we used the ATAC-seq data in Brain Open Chromatin Atlas (BOCA) [123] and ENCODE Consortium for the brain specific (including fetal brain) DNAseI hypersensitive sites [124] to identify chromatin accessible regions (CARs) from 80 kilobases (kb) upstream of the transcription start site (TSS) to the same distance downstream of the 3′ untranslated region (3′ UTR) in the genes of interest. We placed emphasis on the BOCA data that contain CARs for neuronal and non-neuronal cell types across 14 brain regions [125] to identify SNPs associated with neural-specific regulatory elements. The brain regions and abbreviations are described in the Supporting Information document. The smMIPs were designed to minimally overlap each ∼200 bp DHS or ATAC-seq site with an additional 50 bp of flanking sequences. The resulting 1574 smMIPs and a 55 probe smMIP fingerprinting collection were pooled, tested on a set of control DNAs, and rebalanced, resulting in a final pool of 1569 smMIPs (Supporting Information - Methods and Table S1).

#### Multiplexed next generation sequencing

Libraries were prepared [120, 126] and pooled for sequencing in batches of 384. Each pool was sequenced using standard paired-end (100bp) rapid run chemistry in a single lane on a HiSeq 2500 (Illumina, San Diego, CA). The final batch contained repeats from previous batches. Using a quality control benchmark requiring that each sample have a minimum of 80% of target bases covered with a depth of at least 10, 2040 (92%), 2190 (99%), and 2176 (98%) of the 2209 samples prepared passed the benchmark on chromosomes 6, 12, and 15, respectively. For de-multiplexing, generation of FASTQ files, and annotation of sequence data, we used the same in-house pipeline as for MIP design (see Supporting Information). After QC steps, the sample sizes were 1333, 782 and 46 for the UW, SK and UH samples, respectively, with average genotype completion rates of 98.7%, 99.2% and 98.9%, respectively. Variants were annotated using the 1000 Genomes Project (1KGP) and Ensembl Variant Effect Predictor (VEP) [127].

### Statistical and Bioinformatic Analyses

#### Overview

We carried out association analyses on all MIPs sequence variants and samples that passed QC. A full range of variant frequencies including some low-frequency causal risk variants were considered. Low frequency variants were specifically evaluated because each ROI investigated was first identified through linkage analysis, which is most sensitive to low frequency segregating variation.

#### Ancestry adjustment

Self-reported continental ancestry was available for most samples. Because almost all samples were of European origin, either by self-report or KING estimation (Supporting Information), a simple European/non-European indicator assigned by self-report was used to adjust for ancestry in all analyses as a potential nuisance covariate. SNP-based ancestry estimates conflicted with self-reported ancestry in only six individuals. SNP-based ancestry was used in these individuals because self-reported ancestry may reflect cultural affiliation rather than genetic ancestry [128].

#### Phenotypes and adjustments

Reading-related phenotypes used for our analyses that were directly comparable across the three data sets included word identification (WID) and word attack (WA) from the *Woodcock Reading Mastery Test* (*WRMT*; [129], single word reading efficiency (SWE) and phonological decoding efficiency (PDE) from the *Test of Word Reading Efficiency* (*TOWRE;* [130]). The UW and SK cohorts also included spelling (SP) from the *Wide Range Achievement Test – Revised* (*WRAT3-R;* [131]), and nonword repetition (NWR) from the *Comprehensive Test of Phonological Processing* [132]. For all reading-related phenotypes considered here, lower scores indicate more impairment on the measure. In addition, only the UW and SK cohorts included verbal IQ (VIQ), which was used as a covariate for some analyses. Two different phenotype adjustments were considered: (1) adjustment for age and sex only (UNADJ), and (2) adjustment for age, sex and VIQ (VIQADJ). The second set of adjustments represents traits that can be interpreted as free from VIQ effects but could only be performed on the UW sample and children in the SK sample, because of the availability of VIQ. For brevity, when discussing results, we use the format Trait:Adjustment (*e.g*., SWE:UNADJ and SWE:ADJ) to refer to the phenotype without or with VIQ adjustment.

#### Association testing

Phenotypes were adjusted for covariates within each data set using linear regression, and the residuals were used as the response variable in the association testing. Adjustments were done within data sets both because different editions of tests were used across data sets, and because there were differences in ascertainment. Association testing using GENESIS [133] was done using the combined data sets, with distinct means and variances modeled. Data set-specific variances were used to allow for differences between data sets. Family relationships were accounted for by using the expected pedigree-defined kinship in the covariance matrix via a mixed model. For all SNPs observed in two or more copies in the combined data set, we regressed the phenotype of interest against the dose of the minor allele, yielding a single-variant test. Because SNPs with minor allele frequency (MAF) > 0.01 should result in approximately 40 copies in our dataset, we chose this value as the threshold over which statistical analyses should be reliable. All SNPs, including those with lower MAF, were included in aggregate analyses, with grouping according to their location relative to each candidate gene, defined as 5′ region, 3′ region, exons, and introns. The SKAT-O [134] aggregate test was performed in GENESIS, using weights following a Beta distribution with parameters (1,25) and dependent on the MAF [134]. This choice of weight distribution more heavily weights rare variants, but still allows for a contribution from common variants. The SKAT-O test optimizes power by finding the maximal weighted average of the Burden test (more powerful when most variants are causal and effects are in the same direction) and SKAT test (more powerful when most variants are not causal and effects can be in either direction). Correcting for multiple testing is difficult in these data because there are strong correlations between the phenotypes used, and the focus is on small regions of the genome where we expect substantial linkage disequilibrium (LD). As an approximate correction, we require *p* < 0.01 for significant evidence of association, which corresponds to a *p*-value of 0.05 divided by 5, the number of candidate genes being considered.

#### Haplotype Estimation and Testing

When multiple SNPs in LD with one another achieved *p* <0.01, we used Beagle 5.4 [135] with the 1KGP European reference population (EUR) [136] to obtain phased genotypes, thus providing pairs of phased haplotypes for each subject. For a locus with *n* common haplotypes, we fit *n* additive models for each phenotype, where the *i*th model estimates the dose effect of haplotype *i* relative to the other haplotypes pooled. GENESIS allowed us to correct for relationships by using the pedigree-defined kinship in the covariance matrix of a mixed model. When more than one haplotype in a region showed significant association in a dose-model with a phenotype, we fitted a model including dose effects for all the significant haplotypes and allowed for interaction between them.

#### Annotating Non-coding Variants in CARs

We explored the potential impact of all non-coding variants with MAF > 0.01 and *p* < 0.01 on at least one single-variant association test for any of the UNADJ and VIQADJ phenotypes. We annotated variants using the JASPAR tracks on the UCSC Genome Browser [137] as well as the JASPAR database [138]. We considered four characteristics of non-coding variants that together are suggestive of a regulatory effect: 1) the variant is in the peak signal (∼200bp) in either ATAC-seq [125] or DNAseI-seq brain profiles [139] in any brain region; 2) it overlaps with a known TF motif, as found in JASPAR [138]; 3) the change disrupts a conserved position in the motif, as assessed by the position frequency matrices in JASPAR; and 4) the TF whose motif is disrupted has an open promoter in the same brain region(s) as the variant [140–142].

## RESULTS

### Sample Characteristics

#### Ancestry

In the SK data set, all samples with SNP data (all 251 children) had estimated proportion of EUR ancestry greater than 95%. Therefore, the SK data set (including parents) was assumed to be 100% European in genetic background. For the 532 people with SNPs in the UW data set, 504, 15 and 2 individuals had EUR, AFR and East Asian (EAS) ancestry proportion greater than 95%, respectively. Eleven people were admixed (8 EUR/EAS and 3 EUR/AFR) and were counted as non-Europeans. Considering both self-reported ethnicity and SNP-estimated ancestry, 1208 people in the UW data set were assigned to the European category and 100 people to the non-European category. Twenty-five people categorized as unknown because data were unavailable were dropped from the analysis. The UH cohort had 5 African American individuals and 32 white Hispanic individuals as determined by self-report.

#### Phenotypes

In the tables and text that follow, these abbreviations are used for the tests and the processes they assess: **WID** (*WRMT-R Word Identification* for accuracy of oral reading of real words), **WA** (*WRMT-R Word Attack* for accuracy of oral reading of nonwords), **SP** (*WRAT-3* or *WRAT-R Spelling* for written spelling of orally dictated words), **SWE** (*TOWRE* speed of oral reading of real words), **PDE** (*TOWRE* speed of oral reading of nonwords), **NWR** (*CTOPP* Nonword Repetition NWR for phonological memory). Table 1 shows sample sizes for UNADJ and VIQADJ phenotypes for each dataset and the combined dataset. The variability in sample numbers included in the analyses reflects differences in the phenotyping protocols at the three institutions. VIQ scores were obtained for parents and children in the UW cohort, only for children in the SK cohort, and were not obtained for the UH cohort; therefore, the VIQADJ samples include only the UW and SK cohorts. For SWE and PDE, the UNADJ dataset is substantially larger than the VIQADJ dataset, because VIQ was not available for SK parents. Results presented here in the main text focus on the larger, UNADJ, dataset except when findings are substantially different for VIQADJ.

**Table 1:** Sample size by data set for phenotypes analyzed.

| Sample | Phenotype<sup>1</sup> |  |  |  |  |  |  |  |  |  |  |  |
| --- | --- | --- | --- | --- | --- | --- | --- | --- | --- | --- | --- | --- |
|  | WID |  | WA |  | SP |  | SWE |  | PDE |  | NWR |  |
|  | UN<br>ADJ | VIQ<br>ADJ | UN<br>ADJ | VIQ<br>ADJ | UN<br>ADJ | VIQ<br>ADJ | UN<br>ADJ | VIQ<br>ADJ | UN<br>ADJ | VIQ<br>ADJ | UN<br>ADJ | VIQ<br>ADJ |
| UW | 1315* | 1315* | 1315* | 1315* | 1313* | 1313* | 1305* | 1305* | 1304* | 1304* | 1302* | 1302* |
| SK | 335 | 331 | 336 | 332 | 338 | 338 | 713* | 332 | 711* | 332 | 337 | 333 |
| UH | 34 | 0 | 34 | 0 | 0 | 0 | 34 | 0 | 34 | 0 | 0 | 0 |
| **Total** | **1684** | **1646** | **1685** | **1647** | **1651** | **1651** | **2052** | **1637** | **2049** | **1636** | **1639** | **1635** |

1. Word identification (WID) and word attack (WA) subtests of the *Woodcock Reading Mastery Test*, *WRMT* [129]; spelling (SP) subtest of the *Wide Range Achievement Test – Revised*, *WRAT3-R* [131]; single word reading efficiency (SWE) and phonological decoding efficiency (PDE) subtests of the *Test of Word Reading Efficiency*, *TOWRE* [130]; and non-word repetition (NWR) subtest of the *Comprehensive Test of Phonological Processing* [132]. Minor differences in samples size reflect occasional missing values.
\* Includes scores for parents

Supplemental Tables S2 and S3 contain demographic data for the samples used in the UNADJ and VIQADJ analyses with means and standard deviations for the phenotypes in each group. The average VIQ score in the UW data set is almost a standard deviation higher than in the SK data set, as might be expected from the difference in sample selection between the two samples. This is supported by noting that the average score in the UW data set of children (109.7) is not significantly greater than that expected (106.4) from restricting enrollment to VIQ > 90 in a random sample. All the phenotypes have means around zero because they are the residuals from a linear model. The means are not exactly zero because the adjustments were done on a larger data set than only the genotyped participants. Consideration of the SD column demonstrates that the phenotypes fall into two categories: WID, WA and SP where the pre-adjustment value was a standard score, and SWE, PDE and NWR where the pre-adjustment value was a z-score. This difference is reflected in the magnitude of the effect sizes estimated for phenotypes in each category.

### Association Analyses

Tables 2 and 3 summarize results for single variant testing (for variants with MAF > 0.01 only) and aggregate testing, respectively, over all five candidate regions for both the UNADJ and VIQADJ phenotypes. Below we present detailed results primarily on the UNADJ analysis, as it represents the larger data set. Summary tables for the remaining VIQADJ analyses are provided in Tables S4-S6. Complete analysis results for all individual sequence variants for both the UNADJ and VIQADJ phenotypes are also provided in Tables S7 and S8, respectively.

**Table 2:**
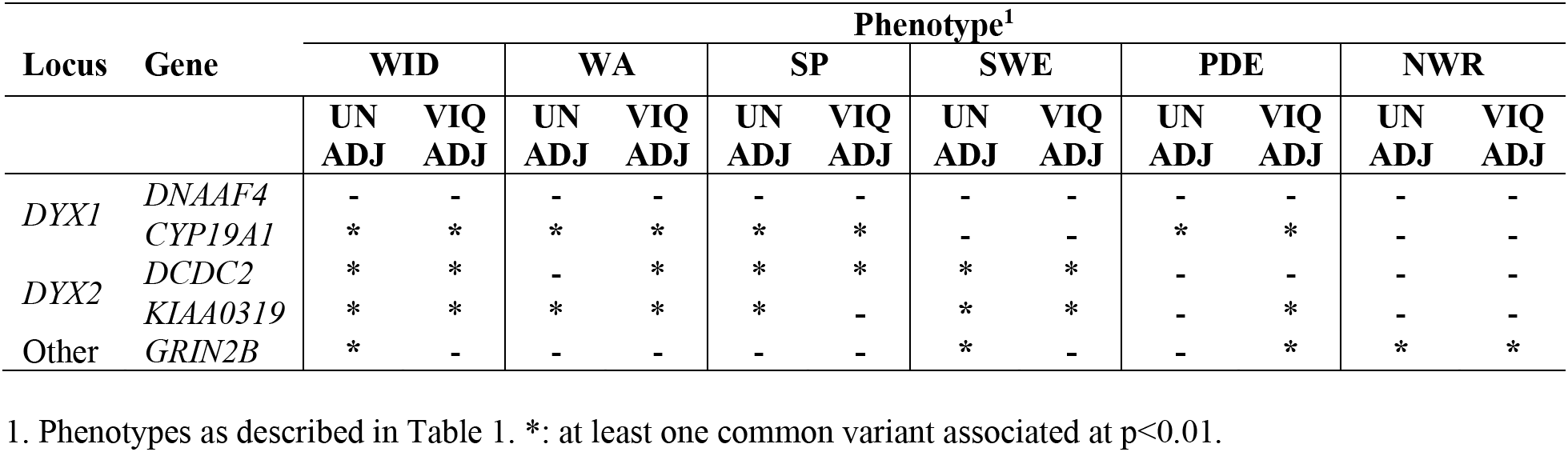
Association (*p*<0.01) of common (MAF > 0.01) variant(s) with UNADJ and VIQADJ phenotypes by candidate gene region.

**Table 3:** SKAT-O p-values for UNADJ, VIQADJ phenotypes, with adjustment for non-European ancestry.

| Locus | Gene<sup>2</sup> | Sites | Phenotype<sup>1</sup> |  |  |  |  |  |  |  |  |  |  |  |
| --- | --- | --- | --- | --- | --- | --- | --- | --- | --- | --- | --- | --- | --- | --- |
|  |  |  | WID |  | WA |  | SP |  | SWE |  | PDE |  | NWR |  |
|  |  |  | UN<br>ADJ | VIQ<br>ADJ | UN<br>ADJ | VIQ<br>ADJ | UN<br>ADJ | VIQ<br>ADJ | UN<br>ADJ | VIQ<br>ADJ | UN<br>ADJ | VIQ<br>ADJ | UN<br>ADJ | VIQ<br>ADJ |
| *DYX1* | *DNAAF4* |  |  |  |  |  |  |  |  |  |  |  |  |  |
|  | DS | 5 | - | - | - | - | - | - | - | - | - | - | - | - |
|  | Exons | 15 | - | - | - | - | - | - | - | - | - | - | - | - |
|  | Introns | 37 | - | - | - | - | - | - | - | - | - | - | - | 0.045 |
|  | US | 20 | - | - | - | - | - | - | - | - | - | - | - | - |
| *DYX1* | *CYP19A1* |  |  |  |  |  |  |  |  |  |  |  |  |  |
|  | DS | 31 | - | - | - | - | - | - | - | - | - | - | - | - |
|  | Exons | 27 | - | - | - | - | - | - | - | - | - | - | - | - |
|  | Introns | 210 | - | 0.027 | - | - | - | - | - | - | - | - | - | - |
|  | US | 105 | - | - | - | - | - | - | - | - | - | - | - | - |
| *DYX2* | *DCDC2* |  |  |  |  |  |  |  |  |  |  |  |  |  |
|  | DS | 34 | - | - | - | - | - | - | - | - | - | - | - | - |
|  | Exons | 54 | - | - | - | - | - | - | - | - | - | - | - | - |
|  | Introns | 112 | - | - | 0.045 | 0.048 | 0.020 | 0.030 | - | - | - | - | - | - |
|  | US | 46 | **0.007** | 0.011 | 0.019 | 0.036 | 0.011 | 0.023 | 0.046 | 0.015 | - | - | - | - |
| *DYX2* | *KIAA0319* |  |  |  |  |  |  |  |  |  |  |  |  |  |
|  | DS | 42 | **0.003** | 0.028 | - | - | 0.031 | - | **0.003** | **0.004** | - | - | - | - |
|  | Exons | 63 | 0.013 | - | **0.007** | 0.026 | **0.007** | 0.036 | - | - | - | - | - | - |
|  | Introns | 145 | - | - | 0.033 | - | - | - | - | - | - | - | - | - |
|  | US | 13 | - | - | - | - | - | - | - | - | - | - | - | - |
| Other | *GRIN2B* |  |  |  |  |  |  |  |  |  |  |  |  |  |
|  | DS | 85 | - | - | - | - | **0.003** | - | - | - | - | - | - | - |
|  | Exons | 50 | - | - | - | - | - | 0.018 | - | - | - | - | - | - |
|  | Introns | 303 | - | - | - | - | 0.024 | **0.008** | - | - | - | - | - | - |
|  | US | 80 | - | - | - | - | - | - | - | - | - | - | - | - |

1. Phenotypes as described in Table 1. **Bold** indicates region/phenotype combinations where $p < 0.01$ . - indicates a $p$ -value $\geq 0.05$ .
2. DS: downstream, US: upstream.

#### DYX1 on chromosome 15

Of the two genes investigated in DYX1, only *CYP19A1* shows evidence of contribution of common variants to the traits analyzed at a significance threshold of *p* < 0.01 whether in the larger UNADJ sample or the smaller VIQADJ sample (Tables 2, S7, and S8). Variants in *CYP19A1* provide significant evidence of association to multiple phenotypes that is robust to presence or absence of adjustment for VIQ (Table 2). In the UNADJ analysis, three SNPs each provide evidence of association with one or more phenotype (Table 4). One SNP, rs55712458, provides highly significant *p*-values for association with four of the six traits, as well as providing the strongest evidence for association across all analyses. This result was robust to phenotypic adjustment, UNADJ vs. VIQADJ (Tables 4 and S4). For three of the four phenotypes with significant evidence of association (WID, WA, and SP), the strength of the evidence is considerably stronger than required to pass our initial screen, particularly in the UNADJ analysis. These results are also stronger than thresholds used in conventional GWAS scans, after factoring in the number of genes evaluated here. A second SNP (rs12594395), located very close to rs55712458, also provides significant evidence for association with WID:UNADJ and has a nominal *p* < 0.05 for WID:VIQADJ (Table S9). Finally, the rare allele of rs141853734 is significantly associated with decreased performance on both WA:UNADJ and WA:VIQADJ.

**Table 4:** Common variants (MAF ≥ 0.01) in *CYP19A1*, UNADJ phenotypes. Effect size (*p*-value) for dose of the rarer allele with adjustment for non-European ancestry. Variants appear in this table if *p* < 0.01 for at least one phenotype.

| rsID | Position<sup>2</sup> | REF/ALT<br>(Freq) | Region<sup>3</sup> | Function<sup>4</sup> | Phenotype<sup>1</sup>: UNADJ |  |  |  |  |  |
| --- | --- | --- | --- | --- | --- | --- | --- | --- | --- | --- |
|  |  |  |  |  | WID | WA | SP | SWE | PDE | NWR |
| rs12594395 | 15:51,483,707 | C/G (0.384) | DS | NA | **1.45 (0.003)** | 1.10 (0.014) | 1.13<br>(0.016) | - | - | - |
| rs55712458 | 15:51,483,996 | G/C (0.198) | DS | NA | **2.82**<br>**(2.7×10<sup>-6</sup>)** | **2.45**<br>**(1.0×10<sup>-5</sup>)** | **2.43**<br>**(2.0×10<sup>-5</sup>)** | 0.10<br>(0.012) | **0.11**<br>**(0.002)** | - |
| rs141853734 | 15:51,627,969 | C/T (0.013) | I | Regulatory | -4.38<br>(0.041) | **-5.36**<br>**(0.006)** | -4.12<br>(0.04) | - | - | - |

1. Phenotypes as described in Table 1. Bold indicates SNP/phenotype combinations where $p < 0.01$ . – indicates $p$ -values $\geq 0.05$ . Freq refers to the alternate allele.
2. Build GRCh37/hg19
3. DS: downstream, I: intronic, E: exonic, US: upstream
4. VEP build38

No aggregate tests in *CYP19A1* or *DNAAF4* yields *p* < 0.01; thus, there is no support for an effect on any of the traits of rare variants in either gene (Table 3). For both genes, only the class of intronic variants provide even suggestive evidence (*p* < 0.05) of association and only for NWR in *DNAAF4* and WID in *CYP19A1*) (Table 3). These aggregate tests were sensitive to the type of phenotypic adjustment, providing, at best, marginal evidence for association only in the VIQADJ and not the UNADJ analysis. Finally, detailed exploration of exonic variants for both genes in DYX1 failed to implicate any variants, common or rare, that could explain more than a tiny fraction of the observed variation in the phenotypes (Tables S7 and S8).

#### DYX2 on chromosome 6

Both common and rare variants in *KIAA0319* show significant evidence for association to multiple phenotypes (Tables 2 and 3). For both common variants and the aggregate tests that weight rare variants more heavily, the phenotypes implicated are primarily WA, WID, SP and SWE, with weaker evidence obtained with PDE and NWR. Two nearly co-located missense variants in *KIAA0319*, both annotated as tolerated/benign, are strongly associated with poor performance on SP:UNADJ (Table 5). In fact, a cluster of four variants (rs2817191, rs2744550, rs2760164 and rs2760163) co-occur in a tiny region of 40 base pairs, and all four variants are strongly associated with a decrease in performance on SP:UNADJ (Table 5). In continental populations of the 1KGP, the variant alleles of these four SNPs are always found together on a single haplotype that occurs with frequency 0.8% in EUR and 15.3% in AFR [143]. The neighboring SNP rs2744549 is in very tight but not complete disequilibrium with the cluster; the slight differences in test statistics and *p*-values for rs2744549 in our data set are explained by missing genotypes in 6 individuals who were typed for the other four variants. The two missense variants (rs2817191 and rs2744550; valine to alanine and serine to glycine, respectively) affect residues that are immediate neighbors in an exon predicted to code a Fibronectin Type III domain [144].

**Table 5:** Common variants (MAF ≥ 0.01) in *KIAA0319*, UNADJ phenotypes. Effect size (*p*-value) for dose of the rarer allele with adjustment for non-European ancestry. Variants appear in this table if *p* < 0.01 for at least one phenotype

| rsID | Position<sup>2</sup> | REF/ALT<br>(Freq)<sup>3</sup> | Region<sup>4</sup> | Function<sup>5</sup> | Phenotype<sup>1</sup>: UNADJ |  |  |  |  |  |
| --- | --- | --- | --- | --- | --- | --- | --- | --- | --- | --- |
|  |  |  |  |  | WID | WA | SP | SWE | PDE | NWR |
| rs114526784 | 6:24,519,996 | T/A<br>(0.053) | DS | NA | - | - | - | **0.22**<br>**(0.003)** | 0.163<br>(0.016) | - |
| rs114979321 | 6:24,544,140 | A/G<br>(0.031) | DS | Regulatory | -2.96<br>(0.028) | - | -3.15<br>(0.021) | **-0.32**<br>**(0.007)** | - | - |
| rs2744549 | 6:24,563,552 | C/T<br>(0.021) | I | NA | **-4.47**<br>**(0.006)** | **-4.14**<br>**(0.005)** | **-4.78**<br>**(0.003)** | -0.24<br>(0.038) | -0.21<br>(0.046) | -0.26<br>(0.024) |
| rs2817191 | 6:24,564,540 | A/G<br>(0.021) | E | missense | -4.00<br>(0.016) | -3.80<br>(0.012) | **-4.25**<br>**(0.009)** | - | - | -0.23<br>(0.049) |
| rs2744550 | 6:24,564,544 | T/C<br>(0.021) | E | missense | -4.00<br>(0.016) | -3.80<br>(0.012) | **-4.25**<br>**(0.009)** | - | - | -0.23<br>(0.049) |
| rs2760164 | 6:24,564,573 | G/A<br>(0.021) | I | splice | -4.00<br>(0.016) | -3.80<br>(0.012) | **-4.25**<br>**(0.009)** | - | - | -0.23<br>(0.049) |
| rs2760163 | 6:24,564,579 | A/G<br>(0.021) | I | splice | -4.00<br>(0.016) | -3.80<br>(0.012) | **-4.25**<br>**(0.009)** | - | - | -0.23<br>(0.049) |

1. Phenotypes as described in Table 1. Bold indicates SNP/phenotype combinations where $p < 0.01$ . – indicates $p$ -values $\geq 0.05$ .
2. Build GRCh37/hg19
3. Freq refers to the alternate allele.
4. DS: downstream, I: intronic, E: exonic, US: upstream
5. VEP build38

The common variants implicated in *KIAA0319* display sensitivity to covariate adjustment and, except for the two missense variants in the common variant cluster, are all in non-coding regions. Aggregate tests also implicate non-coding DNA in the downstream region of *KIAA0319*, and common SNPs with significant association with SWE:UNADJ are in the downstream region (rs114526784 and rs114979321, see Table 5). Finally, there are some variants that only show significant evidence for association with WA:VIQADJ, including rs9295627 and rs9461045 (Table S5). These two SNPs provide nominal evidence for WA:UNADJ (Table S7). SNP rs9461045, here associated with WA:VIQADJ and PDE:VIQADJ (Table S5), was previously shown to affect expression of *KIAA0319* in a different sample [62]. However, WA:UNADJ provides only nominal evidence of association (Table S7), demonstrating moderate sensitivity to the covariate adjustment for this particular SNP. All the SNPs implicated by these analyses are in the large 77kb region that has been previously shown to be associated with reading related phenotypes [53].

Association testing for SNPs in *DCDC2* (Tables 2 and 3) showed sensitivity to covariate adjustment. Table 6 shows two neighboring upstream variants (rs142310124 and rs116652616) significantly associated with SP:UNADJ and SWE:UNADJ, with suggestive evidence for association with WA:UNADJ and WID:UNADJ. In this gene, with adjustment of phenotypes for VIQ, additional intronic variants met criteria for significance for association with SWE:VIQADJ (Table 6). All these additional variants show suggestive evidence for association with SWE:UNADJ at *p* < 0.05 (Table S7). Similarities in minor allele frequencies and estimated effect sizes suggest the presence of at least three haplotype blocks that underlie the region: Block A, rs72830851-rs68182041; Block B, rs12196880 – rs7763196; and Block C, rs77743903 – rs116652616 excluding rs4599626. Block C contains the variants that are also significant with VIQ adjustment (Table 6). Examination of LD among these 9 SNPs in 1KGP EUR [143] shows strong disequilibrium (Dʹ > 0.94) within blocks A, B, and C, and between blocks A and B. The same pattern is seen within 1KGP AFR [143] with the addition of strong disequilibrium between all three blocks. Due to the strong disequilibrium in each block, we cannot statistically distinguish the effects of individual variants; however, bioinformatics prioritization (described below) provides further insights.

**Table 6:** Common variants (MAF ≥ 0.01) in *DCDC2*, UNADJ and VIQADJ phenotypes. Effect size (*p*-value) for dose of the rarer allele with adjustment for non-European ancestry. Variants appear in this table if *p* < 0.01 for at least one phenotype.

| rsID | Position<sup>2</sup> | REF/ALT<br>(Freq)<sup>3</sup> | Region<sup>4</sup> | Function<sup>5</sup> | Phenotype<sup>1</sup>: UNADJ |  |  |  |  |  |
| --- | --- | --- | --- | --- | --- | --- | --- | --- | --- | --- |
|  |  |  |  |  | WID | WA | SP | SWE | PDE | NWR |
| rs77743903 | 6:24,332,778 | A/G<br>(0.021) | I | Regulatory | **-4.74**<br>(0.007) | -3.26<br>(0.042) | -4.03<br>(0.018) | **-0.41**<br>(0.001) | - | - |
| rs142310124* | 6:24,421,582 | A/C<br>(0.023) | US | NA | -4.15<br>(0.010) | -2.95<br>(0.042) | **-4.29**<br>(0.008) | **-0.30**<br>(0.0098) | - | - |
| rs116652616 | 6:24,421,659 | G/A<br>(0.023) | US | NA | -4.15<br>(0.010) | -2.95<br>(0.042) | **-4.29**<br>(0.008) | **-0.30**<br>(0.0098) | - | - |
|  |  |  |  |  | Phenotype<sup>1</sup>: VIQADJ |  |  |  |  |  |
|  |  |  |  |  | WID | WA | SP | SWE | PDE | NWR |
| rs9379643 | 6:24,105,447 | G/A<br>(0.231) | DS | NA | **1.37**<br>(0.006) | - | - | 0.11<br>(0.012) | - | - |
| rs59336318 | 6:24,223,585 | C/T<br>(0.043) | I | Regulatory | - | **-2.61**<br>(0.0099) | - | - | - | -0.15<br>(0.041) |
| rs61232212 | 6:24,223,611 | A/T<br>(0.043) | I | Regulatory | - | **-2.61**<br>(0.0099) | - | - | - | -0.15<br>(0.041) |
| rs72830851 | 6:24,261,168 | A/C<br>(0.105) | I | NA | - | - | - | **0.17**<br>(0.003) | 0.12<br>(0.012) | - |
| rs72830859* | 6:24,269,173 | T/A<br>(0.104) | I | NA | - | - | - | **0.16**<br>(0.004) | 0.12<br>(0.012) | - |
| rs68182041 | 6:24,283,345 | C/T<br>(0.110) | I | Regulatory | - | - | - | **0.17**<br>(0.003) | 0.11<br>(0.024) | - |
| rs12196880 | 6:24,313,747 | C/T<br>(0.092) | I | Regulatory | - | - | -1.51<br>(0.042) | **-0.16**<br>(0.008) | - | - |
| rs12198320* | 6:24,313,828 | G/T<br>(0.092) | I | Regulatory | - | - | -1.50<br>(0.042) | **-0.16**<br>(0.008) | - | - |
| rs7763196* | 6:24,314,065 | A/G<br>(0.092) | I | Regulatory | - | - | -1.48<br>(0.047) | **-0.16**<br>(0.009) | - | - |
| rs77743903 | 6:24,332,778 | A/G<br>(0.021) | I | Regulatory | -3.50<br>(0.018) | - | -3.23<br>(0.036) | **-0.46**<br>(0.0005) | - | - |
| rs4599626 | 6:24,341,357 | C/A<br>(0.242) | I | Regulatory | - | - | - | **0.12**<br>(0.005) | 0.08<br>(0.033) | - |
| rs142310124* | 6:24,421,582 | A/C<br>(0.023) | US | NA | **-4.65**<br>(0.0009) | -3.36<br>(0.017) | **-3.76**<br>(0.0099) | **-0.44**<br>(0.0004) | -0.22<br>(0.041) | - |
| rs116652616 | 6:24,421,659 | G/A<br>(0.023) | US | NA | **-4.65**<br>**(0.0009)** | -3.36<br>(0.017) | **-3.76**<br>**(0.0099)** | **-0.44**<br>**(0.0004)** | -0.22<br>(0.041) | - |

1. Phenotypes as described in Table 1. Bold indicates SNP/phenotype combinations where $p < 0.01$ . – indicates $p$ -values $\geq 0.05$ . \* Alters brain TF binding site
2. Build GRCh37/hg19
3. Freq refers to the alternate allele.
4. DS: downstream, I: intronic, E: exonic, US: upstream
5. VEP build38.

Evaluation of haplotype effects in *DCDC2* provides evidence for more than one important variant in the region. Table 7 shows the six most common haplotypes defined by the nine SNPs in blocks A, B and C, along with haplotype dosage effects as estimated in six separate linear models (Table 7, left). Neither the dosage of the most common haplotype (1: ATC-CGA-AAG) nor haplotypes 3, 5, or 6 are associated with SWE:VIQADJ, whereas haplotypes 2 (CAT-CGA-AAG) and 4 (ATC-TTG-GCA) are associated with significant effects on SWE:VIQADJ that are in opposite directions. Haplotype 2 carries the rarer of two common haplotypes in block A (**CAT** in Table 7) and is associated with an increase in SWE:VIQADJ. Conversely, the rarer of two common haplotypes in block C (**GCA** in Table 7), on haplotype 4, is associated with a decrease in SWE:VIQADJ. The right side of Table 7 shows the final model in which doses of haplotypes 2 and 4 are both included. The estimated effect sizes are very similar to those found in the individual models, and there is no evidence (*p* = 0.16, model not shown) of interaction between haplotypes 2 and 4. These results indicate that the rare haplotype in block A is associated with an increase in SWE:VIQADJ, and the rare haplotype in block C is independently associated with a decrease in SWE:VIQADJ. The region spanned by these haplotypes includes READ1 (a repeat element previously proposed as the causal variant in this region [112–114], which is located between blocks B and C. Due to poor performance of probes targeting READ1 (Figure S1), we were not able to investigate READ1 directly.

**Table 7:** Haplotype models in *DCDC2,* associated with SWE:VIQADJ. The left side shows effect estimates are from six individual haplotype dosage models. The rarer haplotype in each block is in **bold**. The final model on the right shows effect estimates when dose effects are modeled simultaneously.

| Individual haplotype dosage models |  |  |  |  |  | Final model |  |  |
| --- | --- | --- | --- | --- | --- | --- | --- | --- |
| Haplotype/<br>Model | SNP alleles | EUR<br>freq. | Obs.<br>freq. | Effect<br>Estimate | p-value | Term | Effect<br>Estimate | p-value |
| Hap 1 | ATC-CGA-AAG | 0.778 | 0.784 | -0.010 | 0.91 | Intercept | -0.040 | 0.38 |
| Hap 2 | **CAT**-CGA-AAG | 0.109 | 0.105 | 0.174 | 0.002 | Non-Cauc. | -0.19 | 0.11 |
| Hap 3 | ATC-**TTG**-AAG | 0.073 | 0.076 | -0.071 | 0.29 | UW group | 0.022 | 0.70 |
| Hap 4 | ATC-**TTG-GCA** | 0.011 | 0.014 | -0.579 | 0.0001 | Hap 2 dose | 0.17 | 0.004 |
| Hap 5 | ATT-CGA-AAG | 0.004 | 0.006 | 0.044 | 0.84 | Hap 4 dose | -0.56 | 0.0002 |
| Hap 6 | ATC-CGA-ACA | 0.001 | 0.006 | -0.249 | 0.28 |  |  |  |
| Other |  |  | 0.009 |  |  |  |  |  |

Aggregate tests including all variants provide evidence for association in both *KIAA0319* and *DCDC2* on subsets of the traits (Table 3). Variants in *KIAA0319* show evidence for association in aggregate with either downstream or exonic variants for each of the UNADJ real-word phenotypes (*p* = 0.003-0.007), as well as for one VIQADJ real-word phenotype, SWE:VIQADJ (*p* = 0.004). Although aggregate tests do not identify the specific variants of interest, 9 of the 14 variants in the 3′ UTR exonic region of *KIAA0319* have estimated negative contribution to the trait values for both WA:UNADJ and SP:UNADJ (the two phenotypes with significant aggregate test values for exonic variants). Variants upstream of *DCDC2* show significant evidence for association with WID:UNADJ in aggregate testing (p=0.007), and nearly meet our threshold with WID:VIQADJ (*p* = 0.011). Fourteen of the 18 variants in the upstream gene region of *DCDC2* have negative estimated contribution to WID:UNADJ. Like results for single variant testing, evidence for association is usually stronger in the UNADJ phenotypes than the VIQADJ phenotype analyses. However, despite fewer associations that meet the nominal threshold of interest, the same combination of traits and aggregate classes show at least suggestive (*p* < 0.05) association in the VIQADJ phenotypes as in the stronger test results from the UNADJ phenotypes.

#### GRIN2B on chromosome 12

In *GRIN2B*, several common variants show significant, robust evidence of association, primarily with NWR. Four variants in the downstream region and an intronic variant are associated with NWR:UNADJ (Table 8). Three of the downstream variants (rs11055486, rs10772689, and rs4764003) also meet the criterion for association with NWR:VIQADJ (Table S6) and the fourth (rs7307659) gives suggestive evidence of linkage. A single intronic variant (rs61914322) is significantly associated with poorer performance on both WID:UNADJ and SWE:UNADJ, and gives suggestive evidence for WID:VIQADJ and SWE:VIQADJ. Aggregate variant testing in *GRIN2B* shows significant results only for the SP phenotype. Results in Table 3 show significant evidence of association between variants in the downstream region of *GRIN2B* and SP:UNADJ. Variants in intronic regions show significant evidence of association with SP:VIQADJ and provide suggestive evidence of association with SP:UNADJ. There is only suggestive support for a role of variants in the exonic regions for SP:VIQADJ. All such variants are in the 3′ or 5′ UTR regions or lead to synonymous changes, and none provide statistical support for a role on the phenotype.

**Table 8:** Common variants (MAF ≥ 0.01) in *GRIN2B*, UNADJ phenotypes. Effect size (*p*-value) for dose of the rarer allele with adjustment for non-European ancestry. Variants appear in this table if *p* < 0.01 for at least one phenotype.

| rsID | Position<sup>2</sup> | REF/A<br>LT<br>(Freq)<sup>3</sup> | Region<sup>4</sup> | Function<sup>5</sup> | Phenotype<sup>1</sup>: UNADJ |  |  |  |  |  |
| --- | --- | --- | --- | --- | --- | --- | --- | --- | --- | --- |
|  |  |  |  |  | WID | WA | SP | SWE | PDE | NWR |
| rs11055486 | 12:13,656,999 | T/C<br>(0.256) | DS | Regulatory | - | - | - | - | - | **-0.12**<br>**(0.002)** |
| rs10772689 | 12:13,657,684 | T/C<br>(0.529) | DS | Regulatory | - | - | - | - | - | **-0.11**<br>**(0.001)** |
| rs4764003 | 12:13,659,564 | C/T<br>(0.371) | DS | NA | - | - | - | - | - | **0.10**<br>**(0.003)** |
| rs7307659 | 12:13,676,366 | C/T<br>(0.072) | DS | Regulatory | -2.08<br>(0.024) | - | - | - | - | **-0.18**<br>**(0.005)** |
| rs12184498 | 12:13,823,248 | T/C<br>(0.038) | I | NA | -2.56<br>(0.043) | - | - | - | - | **-0.26**<br>**(0.002)** |
| rs61914322 | 12:14,101,470 | G/A<br>(0.100) | I | NA | **-2.26**<br>**(0.005)** | - | - | **-0.14**<br>**(0.008)** | - | - |

1. Phenotypes as described in Table 1. Bold indicates SNP/phenotype combinations where $p < 0.01$ . – indicates $p$ -values $\geq 0.05$ .
2. Build GRCh37/hg19
3. Freq refers to the alternate allele.
4. DS: downstream, I: intronic, E: exonic, US: upstream
5. VEP build38

#### Non-coding variants in CARs

Bioinformatic investigation of JASPAR, an open access database of non-redundant TFs [138] identified ten variants that are within the peak of a CAR and are predicted to disrupt a TF binding site in brain and/or or neuronal tissue (Table 9). These potential regulatory elements are in four of the five candidate genes, adding further support for a possible role in brain development and/or functioning that leads to dyslexia. Of the total 27 non-coding variants identified by association tests (Table S9), 14 fail to meet both requirements of falling in a peak profile identified by ATAC-seq or DNAaseI-seq and overlapping with a known TF motif and were not considered further. The remaining 13 variants reside in CARs and collectively overlap 50 TF motifs. Nineteen of these variant-motif pairs involve a change in a conserved base of the motif, suggesting a likely impact on TF binding (Table 9).

**Table 9:**
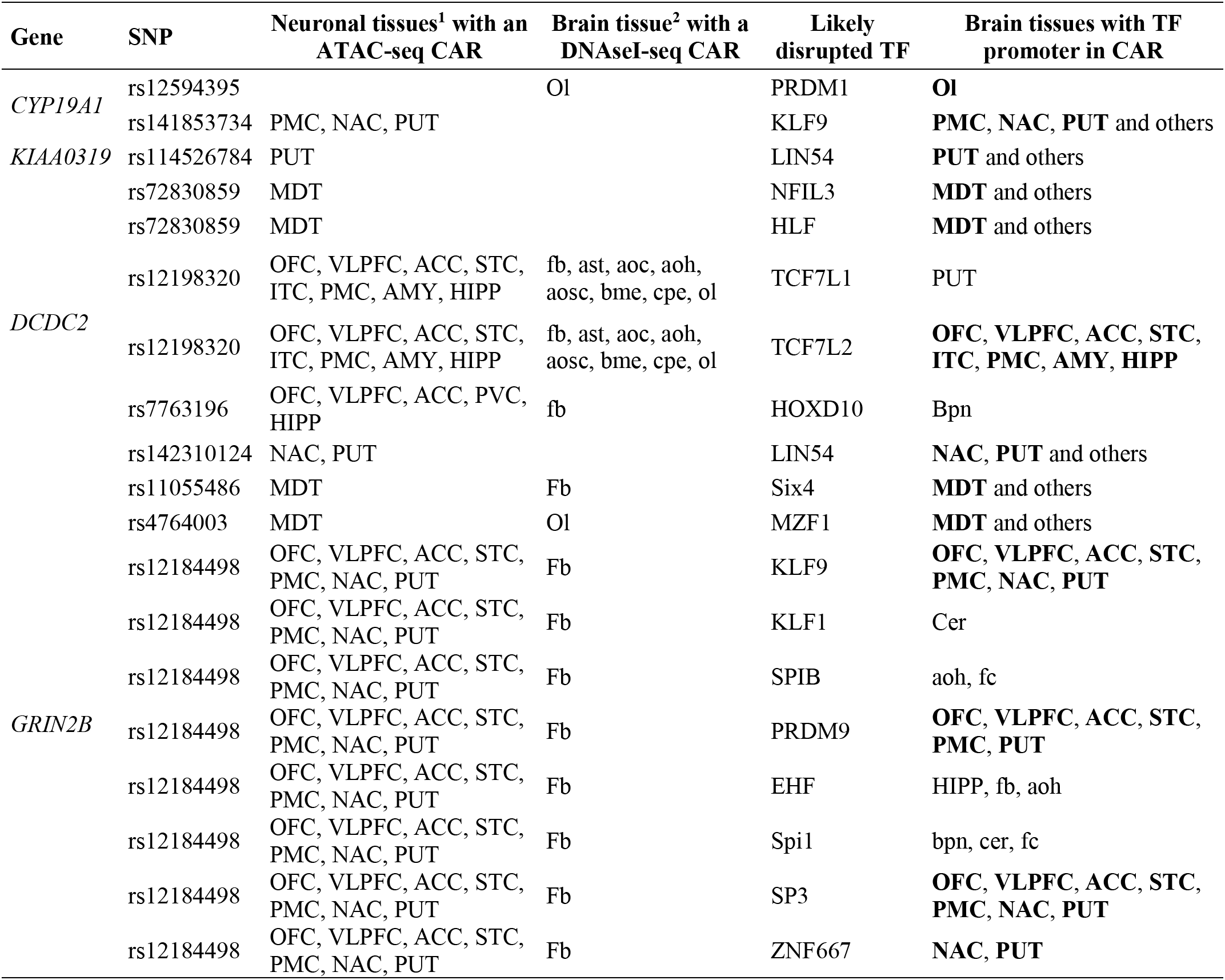
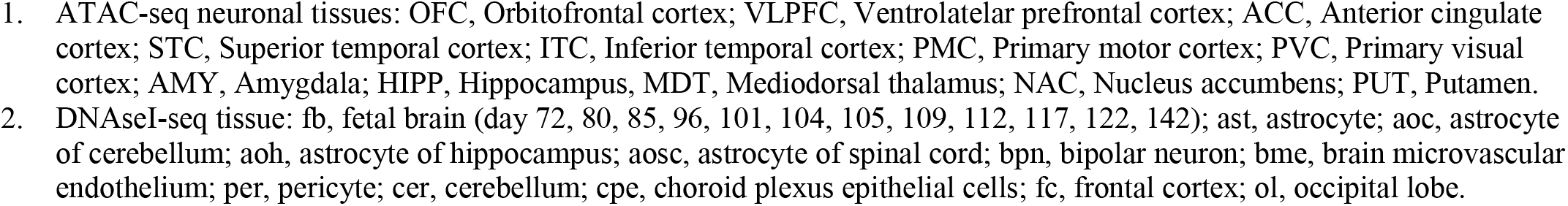
Non-coding variants that are in a CAR and are predicted to disrupt binding of the named transcription factor. Bold in the right-hand column indicates a tissue in which both the SNP and the promotor of the TF are in a <u>CAR.</u>

In *CYP19A1* (Table 4), two of the three non-coding variants potentially have a regulatory role (Table 9). Variant rs12594395 is located 16.1 kb downstream of the 3′UTR in an occipital lobe specific CAR with a predicted loss-of-binding of the TF PRDM1. PRDM1’s promoter is in open chromatin in the same tissue, suggesting it is active in this brain region [140–142] and its function may be disrupted by rs12594395. Variant rs141853734 simultaneously affects two overlapping motifs for TFs KLF9 and ZSCAN4 in neuronal cells of the primary motor cortex (PMC), nucleus accumbens (NAC) and putamen (PUT). KLF9 has an active/poised promoter in these brain regions while ZSCAN4 does not.

In *KIAA0319*, one of two non-coding variants that are not part of the haplotype that carries missense variants (Table 5) may affect TF binding. One (rs114526784) is predicted to disrupt a TF binding site (Table 9). The other (rs114526784) is located on overlapping motifs for TFs ZNF384 and LIN54 in neuronal cells of the PUT. ZNF384 is unperturbed by the nucleotide substitution, but the variant changes the LIN54 motif in a highly conserved fourth residue of the motif that predicts a loss of TF binding. LIN54 has been shown to be a member of the DREAM complex that functions as either a transcription activator or repressor depending on the context [145, 146]. The second SNP, rs11497321, is in a CAR that is present in fetal brain and neuronal cells of 11 adult brain regions, but the resulting nucleotide substitution in NrfI and ELK1:HOXA1 is not predicted to significantly affect binding of either TF.

Several non-coding variants throughout *DCDC2* (Table 6) have the potential for a regulatory role. In the long region defined by the associated haplotype blocks A, B and C described above, there is at least one non-coding variant in each that is predicted to interfere with TF binding. In block A, all three variants are in CARs in some brain tissues, but only rs72830859 in a CAR in mediodorsal thalamus (MDT) results in loss-of-binding mutations predicted to affect TFs PHOX2B, NFIL3 and HLF. Both NFIL3 and HLF have active/poised promoters in all brain regions including MDT. Block B contains a cluster of three variants that span 289 bp and reside within a single CAR that is present in fetal brain and neuronal cells of eight different brain regions. Two of these variants disrupt TF motifs. Variant rs12198320 in Block B is centered within the CAR and disrupts TF motifs for TCF7L1, TCF7L2, and Hnf1A. Hnf1A is inactive in CNS, but TCF7L1 has an active/poised promoter in the neuronal cells of PUT, and TCF7L2 has an active/poised promoter in many neuronal cell types of the brain. Variant rs7763196 in Block B disrupts motifs for TFs CDX4 and HOXD10 in neuronal cells from five brain regions. CDX4 is not expressed in the central nervous system, but HOXD10 has an active/poised promoter in bipolar neurons. Block C variants span 80 kb from the second intron of *DCDC2* to the ninth intron of the downstream gene *MRS2*. One variant, rs142310124, is in a CAR that is specific for neuronal cells in NAC and PUT and is predicted to disrupt motifs for four different TFs (POU2F3, POU3F3, PHOX2B and LIN54). Of these four TFs only LIN54 has an active/poised promoter in PUT. Additional variants rs9379643 and rs61232212 in *DCDC2* are also predicted by JASPAR to disrupt TF binding sites, but neither is present in any neuronal cell types.

Three non-coding variants in *GRIN2B* are predicted to disrupt TF binding. Variants rs11055486 and rs4764003 are both in neuronal-cell specific CARs in MDT and affect the TF motif of Six4 and MZF1, respectively. Both Six4 and MZF1 also have open promoters in MDT. Variant rs12184498 is in CARs present in seven different brain regions and affects eight different TF motifs for KLF9, KLF1, PRDM9, SPIB, EHF, Spi1, SP3 and ZNF667. KLF9, PRDM9, SP3 and ZNF667. All these TFs have open promoters in the relevant tissues, suggesting the availability of these TF for binding. Variant rs10772689 is located within a CAR in some neuronal cells but does not overlap with any known TF motifs. Variant rs7307659 is upstream of a neuronal-specific PUT CAR and was captured in the smMIP experiments, but because it is outside of a peak signal, we are unable to speculate regarding its role.

## DISCUSSION

Here we provided results of a comprehensive investigation of underlying genomic variation in and surrounding five genes with prior evidence for an inherited effect on endophenotypes of dyslexia risk. The MIP sequencing approach that we used allowed inclusion of many more participants and variants than have been previously considered and provided an agnostic approach for identifying underlying causal variants. Reliance of previous studies on detectable linkage disequilibrium between a causal variant and a small number of nearby genotyped polymorphisms is a possible cause of conflicting results across laboratories [51, 52, 72]. In contrast, in the current study, we evaluated much of the gene neighborhoods, focusing on sequence that had the greatest potential for bioinformatic interpretation: protein coding regions and potential regulatory sites upstream and downstream of the candidate genes. Variants evaluated span a wide allele frequency range and fall in both coding and non-coding DNA.

The variants that met our threshold for association and further bioinformatic consideration represent primarily non-coding DNA, with no clearly pathogenic coding variants. Although limited to a small number of selected genes and gene neighborhoods, the results provide an initial prediction of the types of variation that are likely to be more broadly identified through evaluation of DNA sequence in genome-scale studies of specific learning disabilities such as dyslexia. We speculate that variation in coding sequence that results in dramatic alteration of protein structure or function, as is typical of Mendelian disorders, is unlikely to play a role in dyslexia. Such protein-coding variants generally are rare and subject to negative selection, with large impacts on the phenotype. Dyslexia is a phenotype that only is recognized in the presence of widespread literacy, and non-coding variants with subtle effects on gene expression or control are more likely to be relevant. Selection against such variants, with their weak effects on the phenotype, would have been relatively ineffective in the small populations that were typical until very recently in human history. Instead, stochastic effects, such as genetic drift, would have had a role in driving changes in allele frequencies.

Of the variants implicated herein, common variants in *CYP19A1* provide the most significant support for association with all the reading- and spelling-related phenotypes we studied. Bioinformatics evaluation suggests a possible role for regulatory variation that may alter the amount of mRNA, and thus protein, produced. The strongest statistical result was for a variant (rs55712458) in the downstream region of the gene. This variant does not overlap any TF motifs, but it is less than 300 bp away from rs12594395, which occupies an occipital lobe specific CAR, and is expected to affect the binding of the PRDM1 TF. Another variant in *CYP19A1* affects a motif for KLF9, which is a neuronal TF with an active/poised promoter in all brain regions, including fetal brain. In addition to these individual-SNP results, aggregate testing suggests that some probably rare, intronic variants may also be involved. Finally, *CYP19A1* was also previously nominated as a dyslexia-risk gene through identification of the breakpoint of a t(2;15)(p12;q21) translocation that disrupted the promoter region of the gene in a person with dyslexia [37]. *CYP19A1* encodes the estrogen synthase CP19A that converts C19 androgens to C18 estrogens and is responsible for local synthesis of estrogens outside of the reproductive system. In the brain it is expressed from prenatal stages to adulthood [147] in multiple cell types where it regulates synaptic plasticity and plays a role in cognition, memory and language, and many other functions [148, 149].

We also obtained support for involvement in multiple measures of reading/spelling for the two genes evaluated in DYX2: *KIAA0319* and *DCDC2.* In *KIAA0319*, a haplotype that includes missense variants in a predicted fibronectin type III domain [144] is associated with several dyslexia phenotypes. Fibronectin is a protein that has important roles in cell adhesion, migration, differentiation, and proliferation, which has implications for development, among other functions [150–152]. Additional associations were found for common variants, including one (rs114526784) predicted to disrupt a binding site of TF LIN54 in neuronal cells of PUT. For *DCDC2* we detected common intronic variants and both protective and deleterious haplotypes that are significantly associated with dyslexia phenotypes. In particular, two common haplotypes affected timed real word reading scores in opposite directions. Variants within these and other haplotypes are predicted to disrupt binding sites for several TFs. For both genes, therefore, as for *CYP19A1*, some associated variants implicate transcriptional regulation as a possible mechanism of modulation of dyslexia risk.

Finally, common variants downstream of *GRIN2B* are associated with decreased performance on phonological non-word memory, the only phenotype on which this gene had an effect. None of the other studied genes showed an association with this phenotype. One *GRIN2B* variant affects the TF motif of a homeobox protein that may have a role in the differentiation, maturation, and survival of neuronal cells [153–155]. *GRIN2B* was selected for the current study because it lies in a region with evidence of linkage for a phonological non-word memory phenotype in the UW cohort [49] and was associated with verbal memory phenotypes in two European dyslexia datasets [117, 118]. *GRIN2B* encodes GluN2B, one of the glutamate-binding subunits of the tetrameric N-methyl-D-aspartate ionotropic glutamate receptors (NMDARs) that are important for neuronal development and plasticity [156]. GluN2B is highly expressed prenatally in the brain where it is involved in learning and working memory via its role in synaptic plasticity and enhanced long-term potentiation [157, 158]. Pathogenic coding variants and deletions in *GRIN2B* also cause a spectrum of neurodevelopmental disorders [159–161]. Non-coding variants in *GRIN2B* have also been associated with short term and working memory, intelligence quotient and cognitive impairments in dyslexia [117, 118] and with other cognitive and behavioral traits [162–164].

Given diverse opinions about the importance of taking VIQ into account, we note that conclusions in the current study were not strongly affected by adjustment, or not, for verbal IQ. If anything, results were slightly stronger in the absence of VIQ adjustment. Two possible explanations for this observation are (1) that IQ-adjusted analysis had lower power to detect association because of a smaller sample size of participants with the necessary IQ data, and/or (2) use of a covariate with measurement error, as is the case for VIQ, leads to regression to the mean [165], with consequent loss of power to detect the effect of the predictor – the SNP in this case.

There are, of course, some limitations to our study. Although we were able to generate more sequence data on a larger sample than has previously been evaluated for dyslexia, the tradeoff was that the sequencing strategy was limited to a subset of short DNA segments within a small number of previously implicated regions. We therefore cannot comment on genes and genomic regions that fall outside of the regions investigated or sequence alterations such as structural rearrangements or copy number variants (*e.g.*, the READ1 polymorphism [112–114]), that would likely be missed by this short-read technique. Our data in the regions investigated allowed evaluation of most DNA positions in the regions for which current understanding of molecular mechanisms allows bioinformatics interpretation about effects on initiation of transcription. Even so, current knowledge about normal human variation in the regulome is still incomplete, and we acknowledge that transcription factor families share binding motifs, making the definitive identification of specific transcription factors difficult. It is also possible that variants in other regulatory motifs, which can be quite distant from the coding portions of the genes, may hold the causative DNA alterations.

In summary, we provide evidence that variants in or near *DCDC2*, *KIAA0319*, *CYP19A1*, and *GRIN2B* play a role in risk for dyslexia. Given the large sample size, these results argue strongly against the causative involvement of large-effect coding variants in any of the five candidate genes. Instead, these results support a potential role in transcriptional regulation that alters the quantity of RNA produced. The non-coding variant/transcription factor/brain-region combinations we identified provide a starting point for future work on the role of non-coding variants in dyslexia. These results also illustrate some of the challenges that the field will face in identifying causal variants that may act through gene regulation rather than alteration of protein sequences. Use of whole-genome sequence (WGS), especially long-read, would capture regulatory elements with fewer complications, including detection of alterations in repeat sequences that might reside in deep intronic or intergenic regions. However, the WGS approach adds significant cost that could critically limit the number of samples used. The most feasible approach to corroboration of variants and haplotypes of interest discussed herein will therefore require evaluation in other dyslexia sample sets followed by functional studies in an appropriate cell model to begin to determine biological relevance [166], an endeavor that is well beyond the scope of the current analysis.

## ACKNOWLEDGEMENTS

We are grateful to the family members who volunteered their time to participate in the research. John Wolff, Hiep Nguyen, and Edith PA Fuerte provided excellent technical, computational, and bioinformatics assistance. We thank the many graduate student assistants who administered the test batteries.

