## Supplemental information for "Targeted analysis of dyslexia-associated regions on chromosomes 6, 12 and 15 in large multigenerational cohorts"

### Supplemental Methods

#### Ascertainment and Evaluation of the University of Washington (UW) Cohort

Proband recruitment, recruitment of probands' multigenerational family members, and evaluation procedures were done under a protocol approved by the Institutional Review Board of the UW and are described elsewhere in detail [1, 2]. Probands were school age children in grades 1-9 with reading and spelling difficulties based on scores from normed measures of reading and spelling achievement and a Verbal IQ (VIQ) test administered by the research team. Probands who qualified their families for participation in the family genetics study had to have a prorated VIQ ≥ 90 (≥25%ile) on the *Wechsler Intelligence Scale for Children—3rd Edition (WISC-3)* [3], and score below the population mean and at least 1 standard deviation below their VIQ on at least two measures of accuracy or rate of single real or nonword reading or accuracy of spelling from dictation. Most probands met these criteria on most or all those measures. Relatives of the probands, ages 6.5 years and older who also consented were administered age-appropriate measures in the same battery of tests. The full data set consists of 2079 individuals in 284 families. Of these, 1347 individuals from 278 families provided quality DNA samples for sequencing. Phenotypic data were available for 96.8% of the 1333 samples that passed quality control (QC) testing. Family sizes ranged from 3 to 51 individuals, with a median family size of 12 in 2-4 generations. Self-reported ethnicities were as follows: non-Hispanic White (90%), Asian (2.1%), Native American (2.0%), African American (1.1%), Hispanic (0.8%), and Pacific Islander (0.2%).

#### Ascertainment and Evaluation of the SickKids (SK) Cohort

Details of the ascertainment, assessment, and inclusion/exclusion criteria have been described previously [4, 5]. Written informed consent and/or assent was obtained from all participants under protocols approved by the Hospital for Sick Children and University Health Network Research Ethics Boards. Probands were children aged 6–16 in schools in the greater Toronto area and Southern Ontario who had problems reading and were native English speakers or had at least 5 years in an English-speaking school. Probands and their siblings in the same age range were administered a battery of tests including IQ, reading achievement, spelling, and phonological memory. Designation of affected status required WISC-3 or WISC-4 Verbal and Performance IQ [3, 6] ≥ 80 and a score at least 1.5 SD below the mean on 2 of 3 measures of single real- or non-word reading, or 1 SD below the mean on all 3. The phenotype battery in parents was limited to TOWRE measures of single word reading efficiency and phonological decoding efficiency. Self-reported ancestry was available for 185 (76%) of the families. Of these, 180 (73.5%) reported European or European-Canadian ancestry. The remaining families reported small amounts of indigenous ancestry (3 families, 1.6%), African ancestry (1 individual) and Mexican ancestry (1 individual).

#### smMIP Design and Capture

Briefly, a smMIP is an oligonucleotide that has a common 'backbone' flanked by sequences that are specific to the target. The targeting oligonucleotides in the smMIP bind to the same DNA strand, the gap between them is filled in with polymerase, and the circle is closed by ligation. Treatment with exonucleases I and III selectively degrades all linear DNA, leaving only the circularized smMIPs that contain the captured targets of interest. The DNA is amplified to incorporate an adaptor with an 8-base sample-specific index and a 6-base random sequence for molecular tagging. The 8-base index allows multiplex sequencing of up to 384 samples. The 6-base random sequence allows use of informatics to remove sequencing errors [7].

smMIPs were synthesized at 2 μM (Integrated DNA Technologies, Coralville, Iowa), pooled and 5' phosphorylated with 1.25 U of T4 Polynucleotide Kinase (New England Biolabs (NEB), Ipswitch, Massachusetts), 1 X T4 DNA ligase buffer (NEB) and 18.75 μl of smMIP pool (9.5 x $10^{-5}$ pmol/μl) in a 25 μl reaction. Capture reactions combined smMIP pool and 100 ng of genomic DNA at a ratio of 1000 probes per haploid genome. smMIP pool capture performance was assessed using GM12878 DNA (Coriell Institute, Camden, NJ) in a pilot experiment. Individual smMIP were rebalanced for greater performance uniformity among the probe collection. Several smMIP were removed after repeatedly failing to capture the target sequence. To perform the capture, we added 1X FailSafe PCR 2X PreMix B (Illumina, San Diego, CA), 1 U of Ampligase DNA ligase (Lucigen, Middleton, WI), and 1.25 U of FailSafe Enzyme (Lucigen) in a total volume of 10 μl. Reactions were incubated in a thermocycler ($98^{\circ}$C for 5 minutes, followed by $56^{\circ}$C for 5 minutes, lastly $60^{\circ}$C >20 hours). The reactions were chilled on ice while aliquoting exonucleases, 5 U ExoI (NEB), 5 U ExoIII (NEB) and 1X Ampligase buffer (Illumina), then incubated at $37^{\circ}$C for 20 minutes followed by $95^{\circ}$C for 5minutes. The circularized smMIPs containing the captured targets of interest remained post exonuclease digestion. Five microliters of the post ExoI/ExoIII treated DNA was subjected to PCR-amplification with 2X KAPA Library Amplification Kit (Roche, Indianapolis, IN), with 1.6 pmol of a universal forward primer and a unique 9-base index reverse primer (384 unique 9-base barcode per sequencing library) in a volume of 15 μl. PCR reaction condition was $98^{\circ}$C for 5 minutes, then 20 cycles at $98^{\circ}$C for 15 seconds, $60^{\circ}$C for 15 seconds, $72^{\circ}$C for 15 seconds, and a final extension step at $72^{\circ}$C 5 minutes. To assess a successful capture, 3 μl of the PCR reaction was subjected gel electrophoresis analysis to visualize the characteristic 300 bp amplified fragment. In the absence of amplified product, the genomic DNAs were subjected to a second capture experiment. 384 genomic-smMIP capture products were pooled and purified using Agencout AMPure XP beads (Beckman Coulter, Pasadena, CA) following the manufacturer's protocol. A ~300 bp fragment was fractionated on a 2% agarose gel and purified using QIAquick Gel Extraction Kit (Qiagen, Germantown, MD). The DNA is ready for next-generation library preparation.

**Alignment and QC of sequencing reads**

Raw read data were aligned to the genome (hg19) with the Burrows-Wheeler Aligner [8]. Single nucleotide variant and indel calling were performed using the Genome Analysis Toolkit [9]. We excluded variants meeting the following criteria from further analysis: variants with an allele balance > 0.75, quality < 20, quality by depth < 5, or unique capture events < 5. We merged individual VCF files, keeping SNPs that appeared at least twice in the relevant dataset. To distinguish genotypes that are homozygous reference from those that are missing due to a lack of depth at that position, depth for each sample was examined at the position of every variant observed in the data set. If depth was greater than or equal to 10, a genotype not appearing in the VCF file was coded as homozygous reference; otherwise, it was marked missing. Using these criteria, 99.3% of the UW samples had non-missing genotypes for more than 80% of identified variants, followed by 98.0% of the SK samples and 93.8% of the UH samples. Similarly, 93.6% of variants identified in the UW sample had non-missing genotypes for more than 80% of individuals, with corresponding completion rates of 96.3% and 92.5%, respectively, in the SK and UH samples. We removed individuals with > 20% missing data or evidence of family structure errors, and variants with a 20% missing rate.

**Ancestry Estimation**
Existing SNP array genotype data were available on all children in the SK data set, and on 532 individuals in 54 of 278 families in the UW data set. We chose a panel of SNPs in linkage equilibrium and used KING (Chen, 2010) to estimate principal components (PCs) in 4 continental super-populations from the 1000 Genomes Project (AFR, EAS, EUR, SAS). We excluded the AMR super-population because it contains admixed samples from Los Angeles (Mexicans), Puerto Rico, and South America, rather than the North American indigenous populations that we might expect to contribute to our data sets. KING estimates PCs for SNP samples by projection onto the 4-population PC space, and then uses these to estimate ancestry proportions for each sample.

### Supplemental Tables S2 – S6 and Figure S1

Table S2: Demographics of samples, means and standard deviations for UNADJ variables, according to data set.

| | UW Children (407M, 322F) | | | UW Adults (286M, 305F) | | | SK Children (194M, 144F) | | | SK Adults (170M, 211F) | | | UH Children (19M, 16F) | | |
|---|---|---|---|---|---|---|---|---|---|---|---|---|---|---|---|
| | N | Mean | SD | N | Mean | SD | N | Mean | SD | N | Mean | SD | N | Mean | SD |
| **Age(years)** | 729 | 11.7 | 3.38 | 591 | 46.2 | 8.94 | 338 | 10.4 | 2.38 | 381 | 41.5 | 5.64 | 35 | 13.55 | 3.52 |
| **VIQ** | 729 | 109.7 | 12.75 | 590 | 109.0 | 11.67 | 334 | 95.0 | 10.90 | - | - | - | - | - | - |
| **WID** | 727 | 0.142 | 15.47 | 588 | 0.133 | 9.68 | 335 | 0.245 | 13.68 | - | - | - | 34 | -0.045 | 7.87 |
| **WA** | 728 | 0.045 | 12.84 | 587 | -0.074 | 10.66 | 336 | 0.180 | 13.47 | - | - | - | 34 | -0.161 | 6.09 |
| **SP** | 726 | 0.140 | 13.86 | 587 | 0.207 | 12.65 | 338 | 0.150 | 10.45 | - | - | - | - | - | - |
| **SWE** | 719 | 0.011 | 1.24 | 586 | 0.016 | 1.01 | 336 | 0.017 | 0.77 | 377 | -0.003 | 0.79 | 34 | -0.015 | 0.68 |
| **PDE** | 718 | -0.001 | 1.02 | 586 | -0.007 | 0.82 | 336 | 0.012 | 0.73 | 375 | 0.002 | 0.90 | 34 | 0.014 | 0.66 |
| **NWR** | 722 | 0.008 | 0.98 | 580 | 0.002 | 0.87 | 337 | -0.005 | 0.75 | - | - | - | - | - | - |

Table S3: Demographics of samples, and means and standard deviations for VIQADJ variables, according to data set.

| | UW Children (407M, 322F) | | | UW Adults (286M, 305F) | | | SK Children (194M, 440F) | | |
|---|---|---|---|---|---|---|---|---|---|
| | N | Mean | SD | N | Mean | SD | N | Mean | SD |
| **Age(years)** | 729 | 11.7 | 3.38 | 591 | 46.2 | 8.94 | 338 | 10.39 | 2.38 |
| **VIQ** | 729 | 109.7 | 12.75 | 590 | 109.0 | 11.67 | 334 | 95.0 | 10.91 |
| **WID** | 727 | 0.013 | 13.40 | 588 | -0.25 | 7.50 | 331 | 0.24 | 12.9 |
| **WA** | 728 | -0.042 | 11.98 | 587 | -0.35 | 9.40 | 332 | 0.17 | 12.8 |
| **SP** | 726 | 0.033 | 12.87 | 587 | -0.12 | 10.61 | 338 | 0.15 | 10.5 |
| **SWE** | 719 | 0.0 | 1.13 | 586 | -0.010 | 0.90 | 332 | 0.016 | 0.73 |
| **PDE** | 718 | -0.009 | 0.96 | 586 | -0.027 | 0.71 | 332 | 0.012 | 0.70 |
| **NWR** | 722 | 0.002 | 0.90 | 580 | -0.017 | 0.79 | 333 | -0.006 | 0.73 |

Table S4: Common variants (MAF ≥ 0.01) in *CYP19A1*, VIQADJ phenotypes. Effect size (*p*-value) for dose of the rarer allele with adjustment for non-European ancestry. Variants appear in this table if *p*<0.01 for at least one phenotype. Bold indicates SNP/phenotype combinations where *p*<0.01. – indicates a *p*-value ≥ 0.05. Freq refers to the alternate allele.

| rsID | Position[2] | REF/ALT(Freq) | Region[3] | Function[4] | Phenotype[1]: VIQADJ | | | | | |
|---|---|---|---|---|---|---|---|---|---|---|
| | | | | | WID | WA | SP | SWE | PDE | NWR |
| rs55712458 | 15:51,483,996 | G/C (0.198) | DS | NA | **1.99 ($1.3\times10^{-4}$)** | **1.95 ($1.7\times10^{-4}$)** | **2.05 ($1.0\times10^{-4}$)** | 0.10 (0.017) | **0.10 (0.008)** | - |
| rs2899472 | 15:51,516,055 | C/A (0.273) | I | Regulatory | **-1.32 (0.006)** | -1.05 (0.028) | **-1.38 (0.004)** | - | -0.08 (0.022) | - |
| rs141853734 | 15:51,627,969 | C/T (0.013) | I | Regulatory | -4.24 (0.024) | **-4.95 (0.008)** | -3.88 (0.038) | - | - | - |

1) Phenotypes as described in Table 1. 2) Build GRCh37/hg19 3) DS: downstream, I: intronic, E: exonic, US: upstream 4) VEP build38

Table S5: Common variants (MAF ≥ 0.01) in *KIAAA0319*, VIQADJ phenotypes. Effect size (*p*-value) for dose of the rarer allele with adjustment for non-European ancestry. Variants appear in this table if *p*<0.01 for at least one phenotype. Bold indicates SNP/phenotype combinations where *p*<0.01. – indicates a *p*-value ≥ 0.05. Freq refers to the alternate allele.

| rsID | Position[2] | REF/ALT(Freq) | Region[3] | Function[4] | Phenotype[1]: VIQADJ | | | | | |
|---|---|---|---|---|---|---|---|---|---|---|
| | | | | | WID | WA | SP | SWE | PDE | NWR |
| rs114526784 | 6:24,519,996 | T/A (0.053) | DS | NA | - | - | - | **0.22 (0.007)** | - | - |
| rs114979321 | 6:24,544,140 | A/G (0.031) | DS | Regulatory | **-3.24 (0.007)** | **-** | - | **-0.40 (0.0001)** | -0.18 (0.039) | - |
| rs9295627 | 6:24,631,851 | C/T (0.194) | I | Regulatory | - | **1.46 (0.006)** | - | - | 0.08 (0.030) | - |
| rs9461045 | 6:24,649,061 | C/T (0.178) | US | NA | - | **1.52 (0.006)** | - | - | **0.11 (0.008)** | - |

1) Phenotypes as described in Table 1. 2) Build GRCh37/hg19 3) DS: downstream, I: intronic, E: exonic, US: upstream 4) VEP build38

Table S6: Common variants (MAF ≥ 0.01) in *GRIN2B*, VIQADJ phenotypes. Effect size (*p*-value) for dose of the rarer allele with adjustment for non-European ancestry. Variants appear in this table if *p*<0.01 for at least one phenotype. Bold indicates SNP/phenotype combinations where *p*<0.01. – indicates a *p*-value ≥ 0.05.

| **rsID** | **Position[2]** | **REF/ALT(Freq)** | **Region[3]** | **Function[4]** | **Phenotype[1]: VIQADJ** | | | | | |
|---|---|---|---|---|---|---|---|---|---|---|
| | | | | | **WID** | **WA** | **SP** | **SWE** | **PDE** | **NWR** |
| rs11055486 | 12:13,656,999 | T/C (0.256) | DS | Regulatory | - | - | - | - | - | **-0.10 (0.005)** |
| rs10772689 | 12:13,657,684 | T/C (0.529) | DS | Regulatory | - | - | - | - | - | **-0.11 (0.001)** |
| rs4764003 | 12:13,659,564 | C/T (0.371) | DS | Regulatory | - | - | - | 0.08 (0.031) | - | **0.10 (0.002)** |
| rs12184498 | 12:13,823,248 | T/C (0.038) | I | NA | - | - | - | - | **-** | **-0.24 (0.003)** |
| rs2193511 | 12:14,004,787 | G/A (0.370) | I | Regulatory | - | - | - | - | **0.08 (0.007)** | - |

1) Phenotypes as described in Table 1. 2**)** Build GRCh37/hg19 3) DS: downstream, I: intronic, E: exonic, US: upstream 4) VEP build38

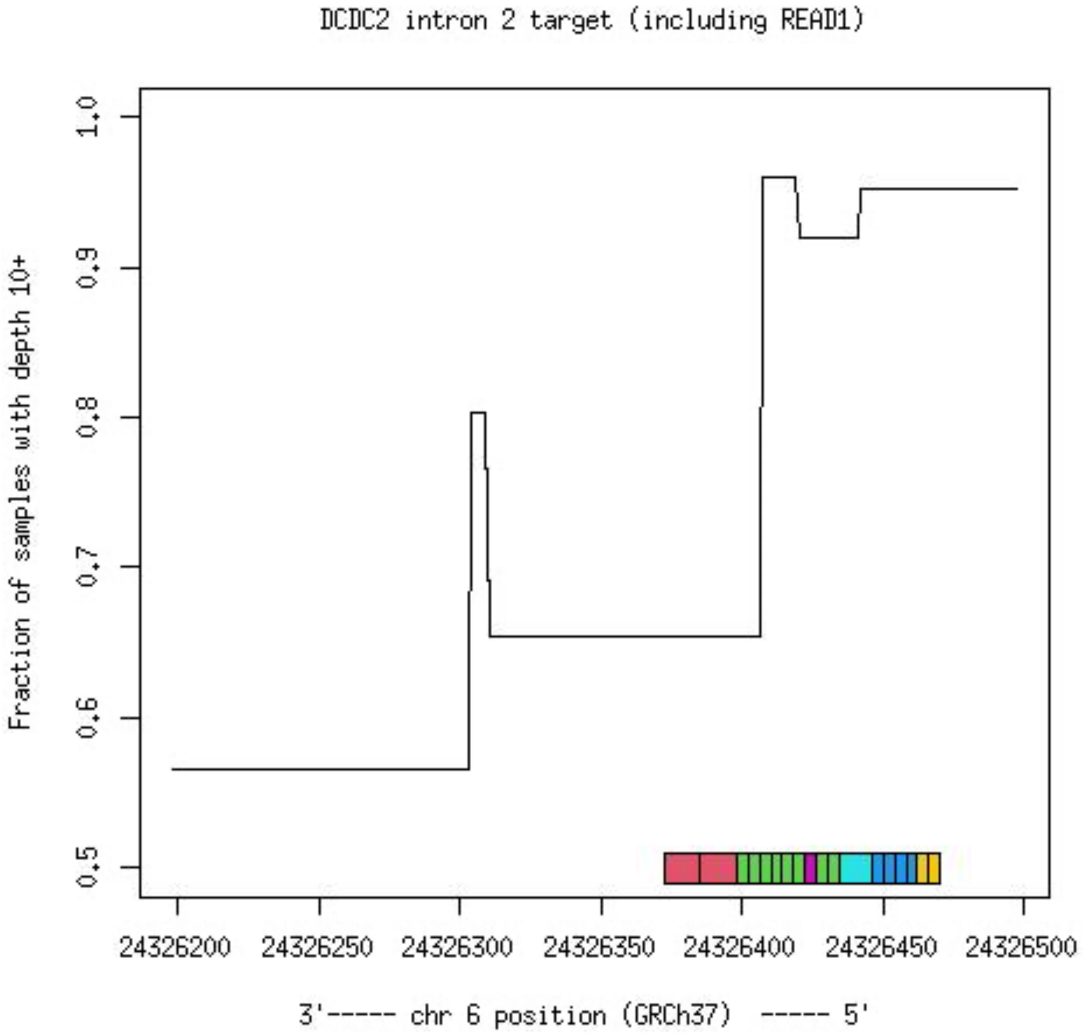


Figure S1: This figure shows the fraction of samples with depth of at least 10 at each position in the target sequence that includes READ1 (found in intron 2 of *DCDC2* on the minus strand). A common deletion that encompasses this entire target is present in ALSPAC at 8.3%. On the 5′ side of the target interval ~ 93% of samples have depth at least 10, consistent with the presence of the deletion in our samples as well. On the 3′ side of the interval only ~57% of samples have depth sufficient for genotype calling. The reference allele of READ1 is depicted by the colored bar. The different colors represent repeated sequences - most of the repeat units are 4 bp long. The risk variants previously associated with dyslexia phenotypes are different numbers of the red repeat unit and first block of green repeat units, in the region where MIP sequence coverage is low. As a result, we are unable to call READ1 alleles in our sample, due to poor performance of our probes on the 3′ end of READ1.
